# Structural recognition of spectinomycin by resistance enzyme ANT(9) from *Enterococcus faecalis*

**DOI:** 10.1101/2020.03.23.003194

**Authors:** Kanchugal P Sandesh, Maria Selmer

## Abstract

Spectinomycin is a ribosome-binding antibiotic that blocks the translocation step of translation. A prevalent resistance mechanism is the modification of the drug by aminoglycoside nucleotidyl transferase (ANT) enzymes of the spectinomycin-specific ANT(9) family or by the dual-specificity ANT(3”)(9) family that also acts on streptomycin. We previously reported the structural mechanism of streptomycin modification by the ANT(3”)(9) AadA from *Salmonella enterica*. ANT(9) from *Enterococcus faecalis* adenylates the 9-hydroxyl of spectinomycin. We here present the first structures of spectinomycin bound to an ANT enzyme. Structures were solved for ANT(9) in apo form, in complex with ATP, spectinomycin and magnesium or in complex with only spectinomycin. ANT(9) shows similar overall structure as AadA with an N-terminal nucleotidyltransferase domain and a C-terminal α-helical domain. Spectinomycin binds close to the entrance of the interdomain cleft, while ATP is buried at the bottom. Upon drug binding, the C-terminal domain rotates by 14 degrees to close the cleft, allowing contacts of both domains with the drug. Comparison with AadA shows that spectinomycin specificity is explained by a straight α_5_ helix and a shorter α_5_-α_6_ loop that would clash with the larger streptomycin substrate. In the active site, we observe two magnesium ions, one of them in a previously un-observed position that may activate the 9-hydroxyl for deprotonation by the catalytic base Glu-86. The observed binding mode for spectinomycin suggests that also spectinamides and aminomethyl spectinomycins, recent spectinomycin analogues with expansions in position 4 of the C ring, will be subjected to modification by ANT(9) and ANT(3”)(9) enzymes.

## INTRODUCTION

Spectinomycin is a broad-spectrum antibiotic that belongs to the aminocyclitol class and was first isolated from *Streptomyces spectabilis* in 1961 (1). It is included in the WHO model list of essential medicines, because of its clinical use to treat gonorrheal infections (2) and is considered to have limited side effects. Spectinomycin is an inhibitor of bacterial protein synthesis. Structural and biochemical studies show that the drug binds to 16S rRNA helix 34 of the 30S subunit of the bacterial ribosome where it blocks the translocation step of protein synthesis (3–5). Spectinomycin is usually bacteriostatic but can be bactericidal at high concentrations (6).

Bacteria have evolved three mechanisms of resistance to spectinomycin. The first is to lower the drug concentration with efflux transporters. This makes some bacteria, for example *Mycobacterium tuberculosis*, intrinsically resistant to spectinomycin (7). The second is alteration of the drug-binding site through chromosomal mutations in the ribosomal 16S rRNA genes (8) or in the gene for ribosomal protein S5 (9). The third is modification of the drug by aminoglycoside modifying enzymes (10–12). Two classes of enzymes modify spectinomycin using ATP, aminoglycoside O-phosphotransferases of the APH(9) class that phosphorylate the antibiotic and aminoglycoside O-nucleotidyl transferases of the ANT(9) and ANT(3”)(9) classes that instead adenylate the antibiotic.

Although spectinomycin is chemically dissimilar to the aminoglycoside streptomycin, both drugs can be adenylated by enzymes of the ANT(3”)(9) (also called ANT(3”)-Ia) class. Aiming to show how one of these, AadA from *Salmonella enterica* (13), recognizes the two chemically distinct molecules in the same active site, we previously solved crystal structures of AadA in its apo form and in complex with ATP, magnesium and streptomycin (14, 15). We could explain how the enzyme coordinates ATP and streptomycin but failed to obtain a structure of AadA with spectinomycin. Furthermore, site-directed mutagenesis followed by *in vivo* resistance measurements and *in vitro* binding assays were used to demonstrate that AadA makes use of different parts of the active site to recognize its two drug substrates. A spectinomycin-binding site was proposed based on manual docking and molecular dynamics simulations. Based on structure-guided sequence analysis, we could confirm that ANT(9) enzymes, which are only active on spectinomycin, are likely to display a similar overall structure. We could also identify sequence determinants that distinguish them from the ANT(3”)(9) enzymes that are active on both antibiotics (15).

Spectinomycin-specific ANT(9) enzymes have been experimentally verified in *Enterococcus faecalis, Staphylococcus aureus and Campylobacter jejuni* and classified into the ANT(9)-Ia and ANT(9)-Ib groups (12, 16–18). Introduction of a plasmid encoding ANT(9) from *E. faecalis* into *E. coli* leads to a 10 000-fold rise of the MIC (minimum inhibitory concentration) of spectinomycin from 10 µg/ml to 100 mg/ml. The ANT(9) enzymes show 29-37 % sequence identity to the dual specificity AadA enzyme from *S. enterica*. In this study, we set out to elucidate how ANT(9) enzymes recognize spectinomycin and how they structurally differ from the ANT(3”)(9) enzymes. To this end, we have solved crystal structures of ANT(9)-Ib from the clinical strain *Enterococcus faecalis* LDR55 (16) in its apo form and in complex with spectinomycin (ANT(9)-spc) or ATP and spectinomycin (ANT(9)-ATP-spc). This structural information allowed detailed analysis of: i) how spectinomycin is recognized by the enzyme, and ii) the structural difference between the single- and dual-specificity enzymes of the same family. In the light of recent developments of novel spectinomycin analogues (19–21), it is most important to understand the molecular basis of the present resistance mechanisms.

## MATERIALS AND METHODS

### Protein expression and purification

The codon-optimized genes encoding ANT(9) from *E. faecalis* (UniProtKB-Q07448) and with a C-terminal His6-tag were synthesized by GenScript and cloned into the the pET11a vector. Plasmids were transformed into *Escherichia.coli* strain BL21 DE3*. A single colony was used to inoculate a five-ml overnight culture of Luria-Bertani (LB) media supplemented with 100 µg/ml ampicillin. The overnight culture was then added to a 1-liter culture and grown at 37 °C until an OD_600_ of 0.5. Protein expression was induced with 1 mM isopropyl-β-thiogalactopyranoside (IPTG) and the culture incubated at 18°C for 20 h. The cells were harvested by centrifugation, washed with 25 mM Tris-HCl pH 8.0, 150 mM NaCl and stored at −20°C.

All the protein purification steps were performed at 8°C. The cells were resuspended in buffer A (50 mM Tris-HCl pH 8.5, 500 mM NaCl, 50 mM each of glutamic acid and arginine and 5 mM β-mercaptoethanol) and 10 mM imidazole and one mini-Complete protease inhibitor tablet, then lysed using a cell disruptor (Constant Systems Ltd., United Kingdom). The lysate was cleared by centrifugation (23,500 × *g*) for 45 min at 4°C. The supernatant was loaded into a gravity-flow column containing Ni-Sepharose High Performance (GE Healthcare, Sweden) pre-equilibrated with buffer A. After incubation for 45 min, the column was washed with 50 ml of buffer A followed by 100 ml of buffer A with 40 mM imidazole. Finally, the protein was eluted with buffer A containing 500 mM imidazole. This fraction was loaded onto a HiLoad 16/60 Superdex 75 prep grade column (GE Healthcare, Sweden) equilibrated with buffer B (25 mM Tris-HCl pH 8.5, 200 mM NaCl, 50 mM each of glutamic acid and arginine) with 5 mM β-mercaptoethanol. The peak fractions containing ANT(9) were pooled and concentrated to 6 mg/ml using a Vivaspin 10 kDa cut-off concentrator (Sartorius AG, Germany).

### Crystallization and data collection

Crystallization trials were performed at 20°C and 8°C by sitting-drop vapor diffusion set up with a Mosquito crystallization robot (TTP Labtech, UK) in 200 nl drops. The sitting drops consisted of a 1:1 ratio of reservoir solution to protein, in which the reservoir solutions were the Morpheus HT-96 screen (Molecular Dimensions, UK). Hexagonal prism-shaped crystals of ANT(9) appeared at 8°C in 10% w/v PEG 4000, 20% v/v glycerol, aminoacids (0.02 M each of sodium L-glutamate, DL-alanine, glycine, DL-lysine, DL-serine), 0.1 M bicine/Trizma base pH 8.5. These crystals were harvested directly from the drop and vitrified in liquid nitrogen. All X-ray data collection was done at 100 K at Diamond Light Source, UK.

Soaking was performed to obtain structures of ANT(9) in complex with ligands. For ANT(9)-spc, crystals were grown in 10% w/v PEG 4000, 20% v/v glycerol, alcohols (0.02 M of each of 1,6-hexanediol, 1-butanol, 0.2 M (RS)-1,2-propanediol, 2-propanol, 1,4-butanediol, 1,3-propanediol), 0.1 M bicine/Trizma base pH 8.5. For the ANT(9)-ATP-spc complex, crystals were grown in 10% w/v PEG 4000, 20% v/v glycerol, monosaccharides (0.02 M of each of D-glucose, D-mannose, D-galactose, L-fucose, D-xylose, N-acetyl-D-glucosamine), 0.1 M MOPS/HEPES-Na pH 7.5. Soaking was done in a 10-µl drop of mother liquor including 10 mM ATP, 10 mM magnesium chloride and spectinomycin powder to saturation. The crystals were soaked for 30 seconds for ANT(9)-spc and 180 seconds for ANT(9)-ATP-spc before vitrification.

### Structure determination and refinement

All diffraction data were scaled and merged using XDS (22) and AIMLESS (23). Xtriage (24) was used to check the data quality and determine the Matthews coefficient. The CCP4 online pipeline MoRDa (25) was used to find a suitable search model and succeeded with PDB ID 5G4A (AadA in complex with ATP and magnesium (15)). The structure was solved by molecular replacement with Phaser (26). The output model from Phaser showed amino acids 8–154 and 166–255 and PHENIX AutoBuild (27) was used for completion of the missing regions between amino acids 154–166. Composite omit maps were calculated for the entire model to check model bias. Manual building was done in COOT (28) and refinement with phenix.refine (29). Protein geometry was validated in MolProbity (30). The ligand bound structures were solved by molecular replacement with Phaser (McCoy et al., 2007) using the apo-ANT(9) structure as template. The polder omit map (31) was calculated to confirm the densities for ATP and spectinomycin.

### Isothermal titration calorimetry

Binding studies were performed at 25°C using a MicroCal iTC200 instrument (GE Healthcare). ANT(9) was used right after the elution from size-exclusion chromatography in buffer B with 5% glycerol and 1 mM tris(2-carboxyethyl)phosphine. Titration was done with 450-650 µM ATP and 200-1000 µM spectinomycin. All ligands were freshly dissolved in the buffer prior to each experiment. For titration of spectinomycin in the presence of ATP, a first titration of ANT(9) with ATP to saturation was followed by a second titration with spectinomycin. The data were analyzed using the MicroCal Analysis plugin in Origin.

## RESULTS AND DISCUSSION

### Protein production and crystallization

ANT(9) from *E. faecalis* was expressed in *E. coli* with a C-terminal His-tag. After size exclusion chromatography, the protein was gradually precipitating. Hence the protein was directly flash frozen in liquid nitrogen for storage or subjected to crystallization. Crystals appeared in a few hours at several crystallization conditions. The apo crystals of ANT(9) diffracted to 1.9 Å and belonged to space group I4 with one molecule in the asymmetric unit. The structure was solved by molecular replacement.

### Overall structure of ANT(9)

ANT(9) has a bi-lobed two-domain structure with approximate dimensions of 52 × 40 × 30 Å^3^ (Figure 1A). The N-terminal domain is a nucleotidyltransferase domain composed of a central five-stranded β-sheet (β_1_- β_5_) and four α-helices (α _1_, α_2_, α_3_ and α_4_). The C-terminal domain is a α-helical domain consisting of five α-helices (α_5_- α_9_). A search for similar structures in the PDB using the DALI server (32) confirmed that AadA from *S. enterica*, which was used as molecular replacement model, was the most similar available structure.

**Figure 1.**
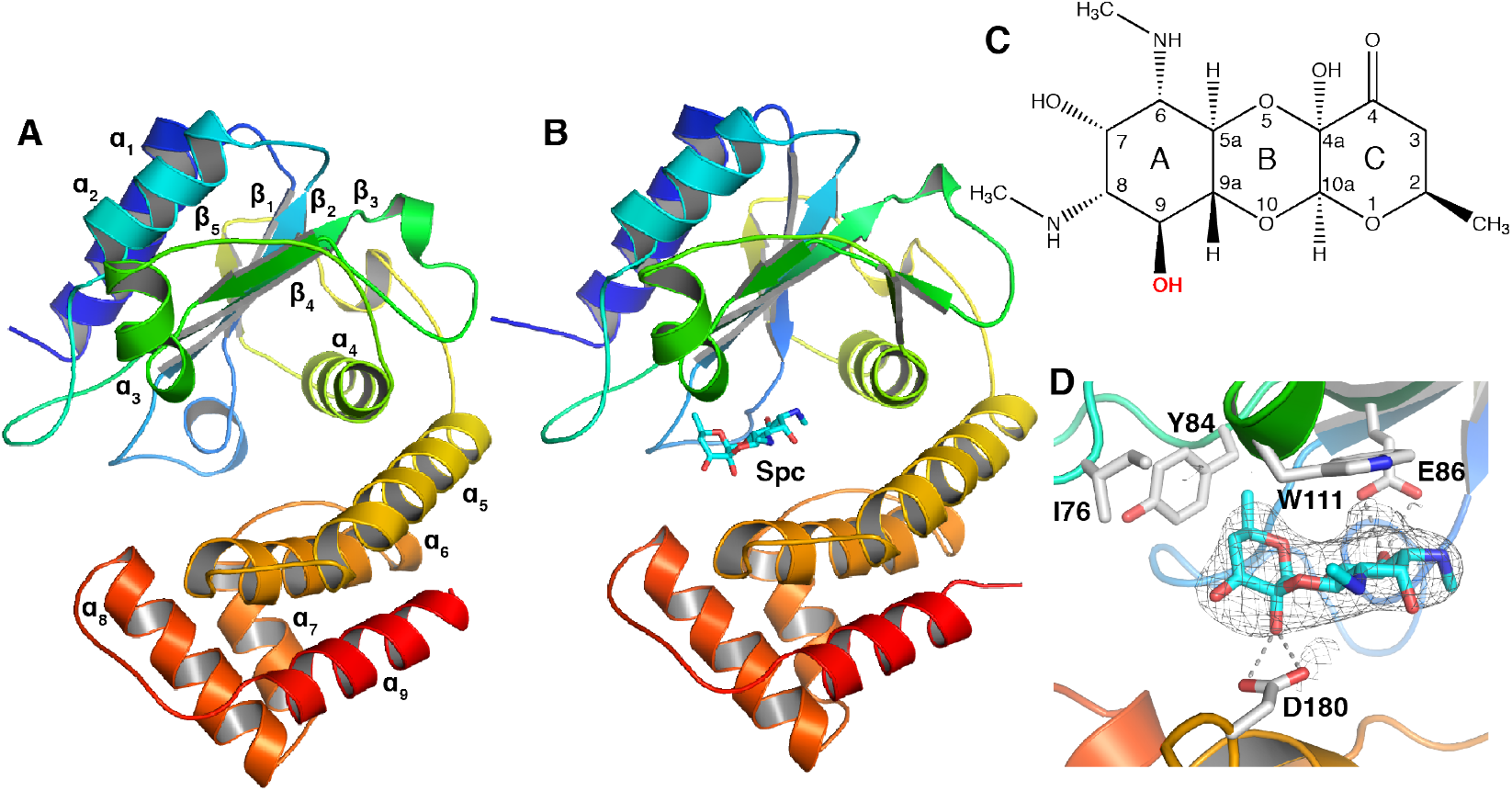
Structure of ANT(9). **A**. Apo structure of ANT(9) represented in rainbow colors going from the N-terminus in blue to the C-terminus in red. **B**. Structure of ANT(9) in complex with spectinomycin (cyan) bound at the entry of the inter-domain cleft. **C**. Chemical structure of spectinomycin. The 9-OH modification site is highlighted in bold. **D**. Detailed interactions of spectinomycin (cyan) in the ANT(9)-spc structure. The F_o_-F_c_ omit map of spectinomycin, contoured at 2.5σ is shown as grey mesh.

### Structures of ANT(9) with spectinomycin and ATP

Attempts to co-crystallize ANT(9) with the substrates ATP and spectinomycin in presence of magnesium failed. Therefore, pre-grown ANT(9) crystals were soaked with ligands, which allowed two structures to be solved; one with only spectinomycin at pH 8.5 and the other with spectinomycin, ATP and Mg at pH 7.5.

The ANT(9) structure with spectinomycin (ANT(9)-spc, Figure 1B) was solved in the same spacegroup (I4) as apo-ANT(9), but with slightly different cell dimensions. The structure was refined to 2.8 Å resolution. We could observe a strong difference density for spectinomycin (Figure 1C-D) but not for ATP or magnesium even though the soak solution contained all ligands.

We suspected that the high pH (8.5) of the crystallization condition might not be optimal for ATP binding. Therefore, further soaking tests were done with crystals grown at pH 7.5. This resulted in a structure of ANT(9) with ATP and spectinomycin (ANT(9)-ATP-spc), for which the space group shifted from I4 to C_121_ with two molecules in the asymmetric unit. The structure was refined to 3 Å resolution, and shows clear electron density for ATP, spectinomycin and one or two magnesium ions in the different chains (Figure 2A, 2B). The two molecules are nearly identical to each other with an RMSD of 0.43 Å over 248 C^α^-atoms.

**Figure 2.**
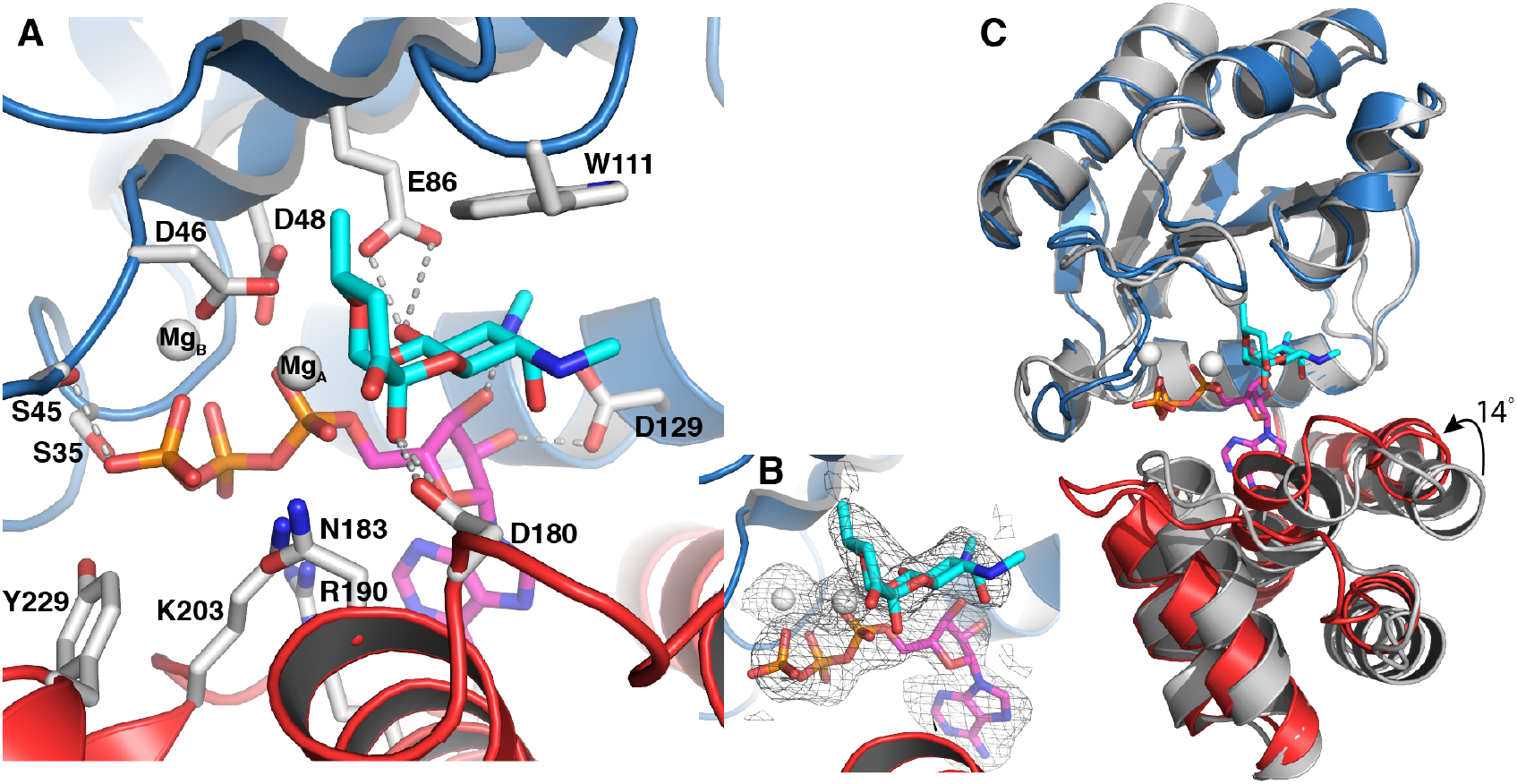
Structure of ANT(9) in complex with spectinomycin (cyan), ATP (magenta) and magnesium (white). The N-terminal domain in blue and the C-terminal domain in red. **A**. Interactions of spectinomycin in the active site of ANT(9)-ATP-spc. Direct hydrogen bonds to spectinomycin are shown as dashed lines. **B.** F_o_-F_c_ omit maps of spectinomycin, ATP and magnesium contoured at 2.5σ. **C**. Superposition of the N-terminal domain of ANT(9) apo (grey) with ANT9-ATP-spc. Upon ligand binding, the C-terminal domain rotates 14° towards the N-terminal domain.

The structures of the individual domains in the ligand complexes are very similar to the apo structure, with an RMSD below 0.8 Å for 150 aligned C^α^-atoms in the N-terminal domain and an RMSD below 0.7 Å over 96 aligned C^α^-atoms in the C-terminal domain. Ligand binding is accommodated by a rotational shift of the second domain towards the first domain (Fig 2C), a shift that also induces changes in cell dimensions and space group. Analysis with DynDom (33) shows that binding of spectinomycin induces a 5-degree rotation of the C-terminal domain. Binding of ATP leads to an additional 9-degree rotation, resulting in up to 4 Å movement at the outer edge of the domain.

In the ANT(9)-spc structure, spectinomycin binds between β_4_ and α_3_ from the N-terminal domain and α_6_ from the C-terminal domain, at the entry of the inter-domain cleft. In the L-shaped spectinomycin molecule, the two first rings are coplanar. The A ring stack against Trp-111 and is coordinated by a hydrogen bond between O9 and Glu-86, the assumed catalytic base (14). The C ring is positioned by a hydrogen bond between the hydroxyl in position 4a and Asp-180 and the insertion of the methyl group in position 2 into a hydrophobic pocket between Ile-76, Tyr-84 and Trp-111 (Figure 1D).

In the ANT(9)-ATP-spc structure, ATP binds at the bottom of the interdomain cleft. In the N-terminal domain, Ser-35, Ser-45, Asp-46 and Asp-48 form hydrogen bonds with the phosphates of ATP, and Asp-129 hydrogen bonds to the ribose. In the C-terminal domain Arg-190, Lys-203 and Tyr-229 form interactions with the ATP phosphates and Arg190 forms a cation-pi interaction with the base (Figure 2A). In both chains, one magnesium ion is coordinated by the three phosphates, Asp46 and Asp-48. In chain A, an additional magnesium ion is coordinated by the alpha phosphate, Asp-46 and O10 of spectinomycin.

In the presence of ATP, spectinomycin together with the C-terminal domain swings 2 Å further into the active site, maintaining the position of the C-ring methyl group. Asp180 is within hydrogen bond distance of both 4a-OH and 4O, while also Asn183 comes within hydrogen bond distance of the 4a-OH. Because of the limited resolution, the exact hydrogen bond network is not clear. In the N-terminal domain the side chain of Glu-86 moves along with the ligand and forms a short or very short (2.7 or 2.2 Å in the two molecules) hydrogen bond with 9O and is within hydrogen bond distance of the secondary amine in position 8. The same amine also forms a hydrogen bond to the 3’ hydroxyl of the ribose of ATP. There are only minor differences between the ligand interactions in the two molecules.

ITC was performed to test the binding affinity of ANT(9) to its substrates and to confirm the specificity to spectinomycin. Due to precipitation problems, these experiments did not generate reproducible K_d_ and stoichiometry values but showed micromolar affinity binding to both ATP and spectinomycin. Despite our observation that spectinomycin could bind in absence of ATP in the crystal structure, we could not observe spectinomycin binding in solution in absence of ATP (data not shown). This suggests that in solution, binding of ATP may position the two domains for interaction with the antibiotic substrate, as previously observed for AadA (15).

### Comparison of ANT(9) with the dual-specificity AadA

Comparison of ANT(9)-ATP-spc with the structure of the dual specificity ANT(3”)(9) AadA in complex with ATP and streptomycin (AadA-ATP-sry) shows that their overall structures are very similar (Figure 3A). The N-terminal domains superpose with an RMSD of 0.95 Å over 110 Cα atoms and the C-terminal domains with an RMSD of 0.83 Å over 62 Cα atoms. The interdomain orientation is slightly different, leading to an RMSD of 1.5 Å over 230 Cα atoms when the complete structures are superposed.

**Figure 3.**
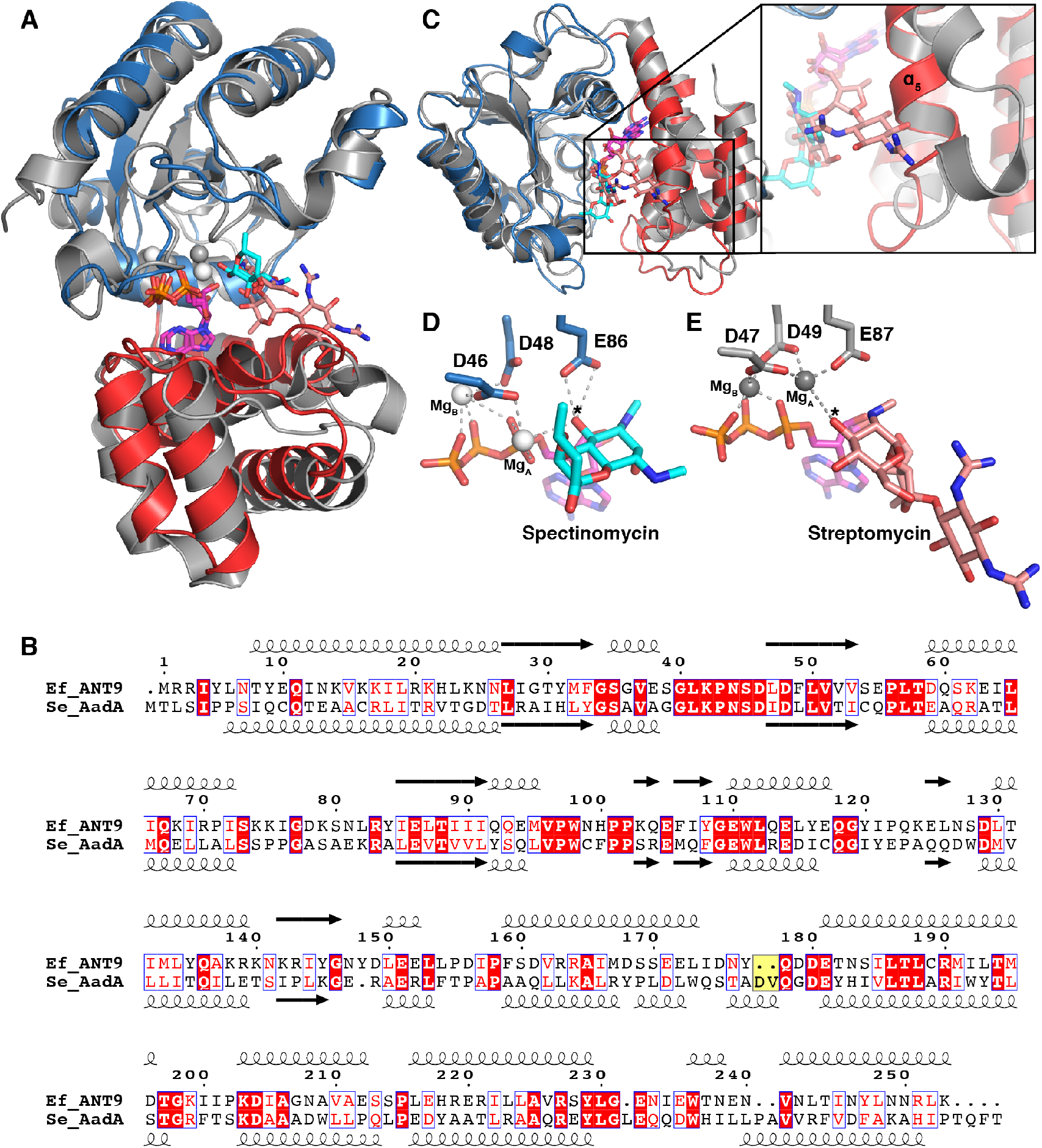
Comparison of ANT(9) and AadA. **A**. Superposition of the N-terminal domain of ANT(9)-ATP-spc (colors as in Figure 2) with AadA in complex with ATP and streptomycin ((15) grey, streptomycin in salmon). **B**. Structure-guided sequence alignment of *E. faecalis* ANT(9) with *S. enterica* AadA (15, 42). The characteristic Asp-Val insertion in AadA is highlighted in yellow. **C.**The straight α_5_ in ANT(9) would clash with streptomycin. AadA shows a kink in helix α_5_ and the insertion in the α_5_ – α_6_ loop forms a short helix to accommodate streptomycin (salmon). View is perpendicular to A. **D-E**. Comparison of active site and magnesium coordination in ANT(9)-ATP-spc (D) and AadA-ATP-str (E). The asterisk indicates the substrate hydroxyl to be modified.

Previous structure-guided sequence alignments showed that the active-site residues that bind to ATP and spectinomycin are conserved between ANT(3”)(9) and ANT(9) enzymes. Enzymes with activity on streptomycin contain a characteristic Asp-Val or Asp-Trp insertion in the loop after α_5_ (Figure 3B). In this region, Trp-173 and Asp-178 were through site-directed mutagenesis shown to be critical for streptomycin binding and resistance, but not for spectinomycin (15).

Looking closer at this region of the superposed structures (Figure 3C), a striking difference is observed for amino acids 160-185 in the α_5_-α_6_ region of the C-terminal domain. In AadA-ATP-sry, Pro169 in helix α_5_ kinks the helix approximately 10 degrees away from the cleft. This, combined with the extension of the loop to form a short helix allows the enzyme to accommodate the large streptomycin substrate. In contrast, the corresponding region of the C-terminal domain in ANT(9)-ATP-spc forms a straight helix α_5_ and a short loop followed by helix α_6_. In the loop, Asp-180 and Asn-183 hydrogen bond to spectinomycin in a position where they would clash with a larger substrate such as streptomycin.

Previous site-directed mutagenesis of Asp-182 in AadA, the equivalent of Asp-180 in ANT(9), showed that the Asp182Asn mutation abolishes binding of spectinomycin to AadA while not affecting binding of ATP or streptomycin. *In vivo*, the Asp182Asn, Asp182Ala and also the Trp112Phe and Trp112Ala mutations are more detrimental for resistance to spectinomycin than streptomycin (15). In the present ANT(9) complex structures, only three amino acids, Glu-86, Asp-180 and Asn-183 (Figure 2A), form direct hydrogen bonds to spectinomycin, thus explaining why Asp180 is critical for spectinomycin resistance. The small and rigid spectinomycin molecule requires closing of the inter-domain cleft to allow snug packing against the N-terminal domain with only two hydrogen bonds to the C-terminal domain. Our analysis predicts that a similar domain closure would need to occur during binding of spectinomycin to AadA and that the binding sites will be identical. Previous MD simulations of AadA with spectinomycin and ATP were not long enough to capture any larger conformational changes of the protein as observed here. The larger streptomycin molecule is instead positioned by stacking with the equivalent of Trp111 and hydrogen bonds to three residues of the C-terminal domain up to 15 Å away from the catalytic base.

The only other structurally characterized resistance-mediating enzyme in complex with spectinomycin is APH(9)-Ia that phosphorylates the same hydroxyl group as ANT(9) adenylates. Compared to ANT(9), APH(9)-Ia forms more extensive hydrogen bond interactions with spectinomycin, at five different positions of the drug (34).

### Catalytic site

Previous *in vivo* MIC experiment and *in vitro* adenylation experiment on AadA demonstrated that the equivalent of Glu-86 in ANT(9) is the catalytic base. Adenyl transferase enzymes, similar to polymerases, make use of a two metal-ion mechanism (15, 35). In ANT(9)-ATP-spc, one magnesium (Mg_B_, Fig. 2A) is similarly positioned as previously observed in AadA and polymerases. Interestingly, the second magnesium (Mg_A_), which displayed clear density in chain A but weak density in chain B, is shifted by 3 Å towards the substrate 9-hydroxyl when compared to in AadA (Figure 3D-E). The close proximity between Mg_A_ and the 9-hydroxyl (3.2 Å) will likely lower the pKa of the 9-hydroxyl to allow the catalytic base Glu-86 to accept its proton. The hydrogen bond between Glu86 and the 9-hydroxyl is very short, 2.2 Å. Computational prediction using H++ (36) suggests that the pKa of Glu-86 is substantially up-shifted in the structural context of the active site allowing its activity in acid-base catalysis at physiological pH. The distance between the 9-oxygen of spectinomycin and the alpha phosphorous of ATP that would be subjected to a nucleophilic attack is 3.9-4.3 Å in our structures (Figure 3D), thus making this structural snapshot closer to the reactive state than previous structures of AadA-ATP-sry (Figure 3E). The active site is organized in agreement with the single in-line displacement mechanism proposed for other adenylyltransferases, KNTase (37) and LinB (38), but the shifted metal ion site may represent a previously unobserved reactive state.

### Conclusions and future outlook

We have here for the first time shown how spectinomycin is recognized by ANT resistance enzymes, and observed an active site arrangement that is closer to the reactive state than what has previously been observed for ANT enzymes. During the late 1980s, there were many attempts to make spectinomycin variants for treatment of bacterial infections (39, 40), but none of these drugs are in clinical use. Lately, however, semi-synthetic modifications to spectinomycin, spectinamides and aminomethyl spectinomycins, have been proposed for treatment of multi-drug resistant tuberculosis and other infections (19, 21). In these molecules, added functional groups on the 4-O position of the C-ring of spectinomycin mediate additional contacts in the spectinomycin-binding site of the ribosome and importantly, prevent the drug from being exported by the intrinsic efflux pump of *M. tuberculosis*. In the APH (9)-Ia, the 4-O position resides within the active site pocket and is involved in water-mediated contacts with the enzyme (34). However in ANT(9), the 4-O is pointing out of the active site. Although promising, these modified drugs will probably still be subject to modification at position 9 by the ANT(9) and ANT(3”)(9) classes of enzymes. To make sure that a new drug does not directly suffer from any of the present resistance mechanisms, it is important to map the contacts of all resistance enzymes on the drug, as previously done for other aminoglycosides (41). In addition, the combination of few critical interactions between ANT(9) and its antibiotic substrate and a substrate-binding site that can be modulated in size based on inter-domain flexibility indicates that this class of resistance enzymes may be prone to evolve in response to other new related antibiotic molecules.

**Table 1.**

|  | ANT(9) | ANT(9)-spc | ANT(9)-ATP-spc |
| --- | --- | --- | --- |
| **Data collection** |  |  |  |
| Beamline | DLS I04-1 | DLS I04 | DLS I04 |
| Wavelength (Å) | 0.915 | 0.979 | 0.979 |
| Space group | I 4 | I 4 | C 1 2 1 |
| Unit cell parameters: |  |  |  |
| a, b, c (Å) | 103.97, 103.97, 61 | 99.6, 99.6, 61.3 | 133.5, 69.3, 94.5 |
| $\alpha, \beta, \gamma$ (°) | 90, 90, 90 | 90, 90, 90 | 90, 135, 90 |
| Resolution (Å)* | 36.8 - 2.1 (2.17 - 2.1) | 49.8 - 2.8 (2.9 - 2.8) | 55.9 - 3.0 (3.1 - 3.0) |
| R<sub>meas</sub>(%)* | 5.8 (161.8) | 14.7(160.3) | 32.4(145.7) |
| <I/σ(I)>* | 25.3 (1.5) | 19.8 (2.2) | 5.4 (1.2) |
| CC ½ (%)* | 100 (79.5) | 99.9(90.4) | 98(47.2) |
| Completeness (%)* | 99.1 (97.3) | 99.57 (99.5) | 99.85 (99.9) |
| Redundancy* | 13.7 (13.8) | 22.9 (23.6) | 6.3 (6.3) |
| **Refinement** |  |  |  |
| Resolution (Å) | 36.98 - 2.10 | 49.8 - 2.8 | 55.9 – 3.0 |
| Reflections / test set | 18972/1847 | 7482/747 | 12341/1186 |
| R<sub>work</sub>/R<sub>free</sub> (%) | 23.0/ 27.2 | 24.3/29.6 | 24/27.1 |
| No. atoms |  |  |  |
| Protein | 2047 | 2047 | 4062 |
| Ligand/ion | 6 | 23 | 113 |
| Water | 16 |  | 12 |
| B-factors |  |  |  |
| Protein | 68.0 | 75.9 | 44.2 |
| Ligands | 78.4 | 71.0 | 38.4 |
| Solvent | 56.0 |  | 32.6 |
| RMSD from ideal |  |  |  |
| bond lengths (Å) | 0.005 | 0.003 | 0.004 |
| bond angles (°) | 0.7 | 0.6 | 0.8 |
| Ramachandran plot: |  |  |  |
| Preferred (%) | 95.6 | 96 | 94.3 |
| Allowed (%) | 4.4 | 4.0 | 5.7 |
| Outliers (%) | 0 | 0 | 0 |
| **PDB accession number** | 6SXJ | 6XZ0 | 6XXQ |

\* Highest resolution shell is shown in parenthesis.

## Acknowledgements

This work was supported by grants 2017-03827 and 2016-06264 from the Swedish Research Council to MS. We are grateful for access to beamlines IO4 and IO4-1 at the Diamond Light Source, Didcot, UK (proposals MX15868 and MX22906). We thank Terese Bergfors for constructive comments on the manuscript.

